# Quantification of Receptor Binding from Response Data Obtained at Different Receptor Levels: A Simple Individual Sigmoid Fitting and a Unified SABRE Approach

**DOI:** 10.1101/2022.06.27.497811

**Authors:** Peter Buchwald

**Author notes:** Correspondence to: Peter Buchwald, Diabetes Research Institute, Miller School of Medicine, University of Miami, Miami, FL 33136, USA.

## Abstract

Methods that allow quantification of receptor binding (occupancy) by measuring response (effect) data only are of interest as they can be used to allow characterization of binding properties (e.g., dissociation constant, *K*_d_) without having to perform explicit ligand binding experiments that require different setups (e.g., use of labeled ligands). However, since response depends not just on the binding affinity-determined receptor occupancy, but also on receptor activation, which is affected by ligand efficacy (plus constitutive activity, if present), and downstream pathway amplification, this requires the acquisition and fitting of multiple concentration-response data. Here, two alternative methods, which both are straightforward to implement using nonlinear regression software, are described to fit such multiple responses measured at different receptor levels that can be obtained, for example, by partial irreversible receptor inactivation (i.e., Furchgott method) or different expression levels. One is a simple method via straightforward fitting of each response with sigmoid functions and estimation of *K*_d_ from the obtained *E*_max_ and EC_50_ values as *K*_d_=(*E*_max_·EC’_50_−*E*’_max_·EC_50_)/(*E*_max_−*E*’_max_). This is less error-prone than the original Furchgott method of double-reciprocal fit and simpler than alternatives that require concentration interpolations, thus, should allow more widespread use of this so-far underutilized approach to estimate binding properties. Relative efficacies can then be compared using *E*_max_·*K*_d_/EC_50_ values. The other is a complex method that uses the SABRE receptor model to obtain a unified fit of the multiple concentration-response curves with a single set of parameters that include binding affinity *K*_d_, efficacy *ε*, amplification *γ*, and Hill coefficient *n*. Illustrations with simulated and experimental data are presented including with activity data of three muscarinic agonists measured in rabbit myocardium.

## INTRODUCTION

Quantitative models linking ligand concentrations, [L], to generated effects, *E*, are fundamental in pharmacology. Not only are they essential in establishing concentration-response relationships, which are the ultimate proof of efficacy and/or safety, but they can also provide valuable insight into the mechanisms of action of ligands and their receptors. Widely used quantitative pharmacology parameters such as the half-maximal effective or inhibitory concentration (EC_50_, IC_50_) are routinely derived from fitting with sigmoid-type equations such as the Hill equation (eq. 1) or its simplified form (*n* = 1, Clark equation) corresponding to a straightforward law of mass action:

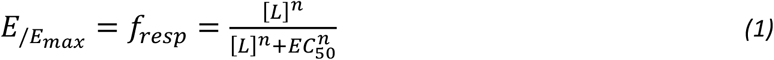

While such functional forms often provide good fit, they may not reveal much about the signaling mechanism or even binding to the receptor because the overall response (effect, *E*) depends not just on receptor occupancy (i.e., the number or fraction of receptors that are ligand-bound), but also on receptor activation (i.e., the number of fraction of receptors that are active either due to the presence of a ligand or to constitutive activity) and on possible downstream pathway amplification, which sometimes can be quite considerable. Thus, the parameter obtained from fitting eq. 1 as corresponding to half-maximal effect (EC_50_, *K*_obs_) is not necessarily closely related to the classic equilibrium dissociation constant *K*_d_ that corresponds to half-maximal occupancy and determines (fractional) receptor occupancy via a similar sigmoid function:

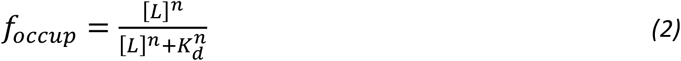

In other words, while both fractional response, *f*_resp_=*E*_/*E*max_, and fractional occupancy, *f*_occup_ = [LR_occup_]/[LR_max_], are typically well described by such equations, they do not overlap and there could be considerable separation between them • Well-known causes of such separations include, for example • partial agonism (where response lags behind occupancy and full occupancy does not result in full response), • signal amplification (where response runs ahead of occupancy and low occupancy can produce full or close to full response creating the appearance of “receptor reserve” or “spare receptors”), • different responses assessed at different downstream vantage points along a signaling pathway (where the same occupancy produces different responses at different readout points), and • biased agonism (where the same occupancy produces different responses along distinct divergent downstream pathways originating from the same receptor) – see [1] for representative illustrations.

Hence, the exact relationship between [L] and *E* (even expressed as normalized fractional response *f*_resp_ = *E*/*E*_max_) is complex, and simultaneous fitting of occupancy and response functions requires models with multiple parameters. For example, our recently introduced SABRE quantitative receptor model provides a framework to fit a variety of response data from the simplest to those of increasing complexity with a unified equation using a total of five parameters in its most general form: three characterizing the ligand-receptor interaction (binding affinity, *K*_d_, efficacy, *ε*, and Hill coefficient, *n*) and two the receptor and its signaling pathway (constitutive activity, *ε*_R0_, and amplification, *γ*) (Figure 1; eq. 17 in Methods and Models) [1-3]. In case of no constitutive activity (*ε*_R0_ = 0), this reduces to the form that will be used here (eq. 18 in Methods and Models), which can also be rearranged to resemble that of eq. 1 above:

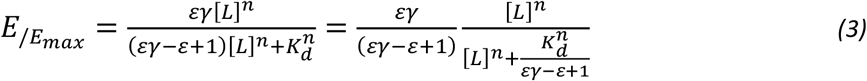

**Figure 1.**
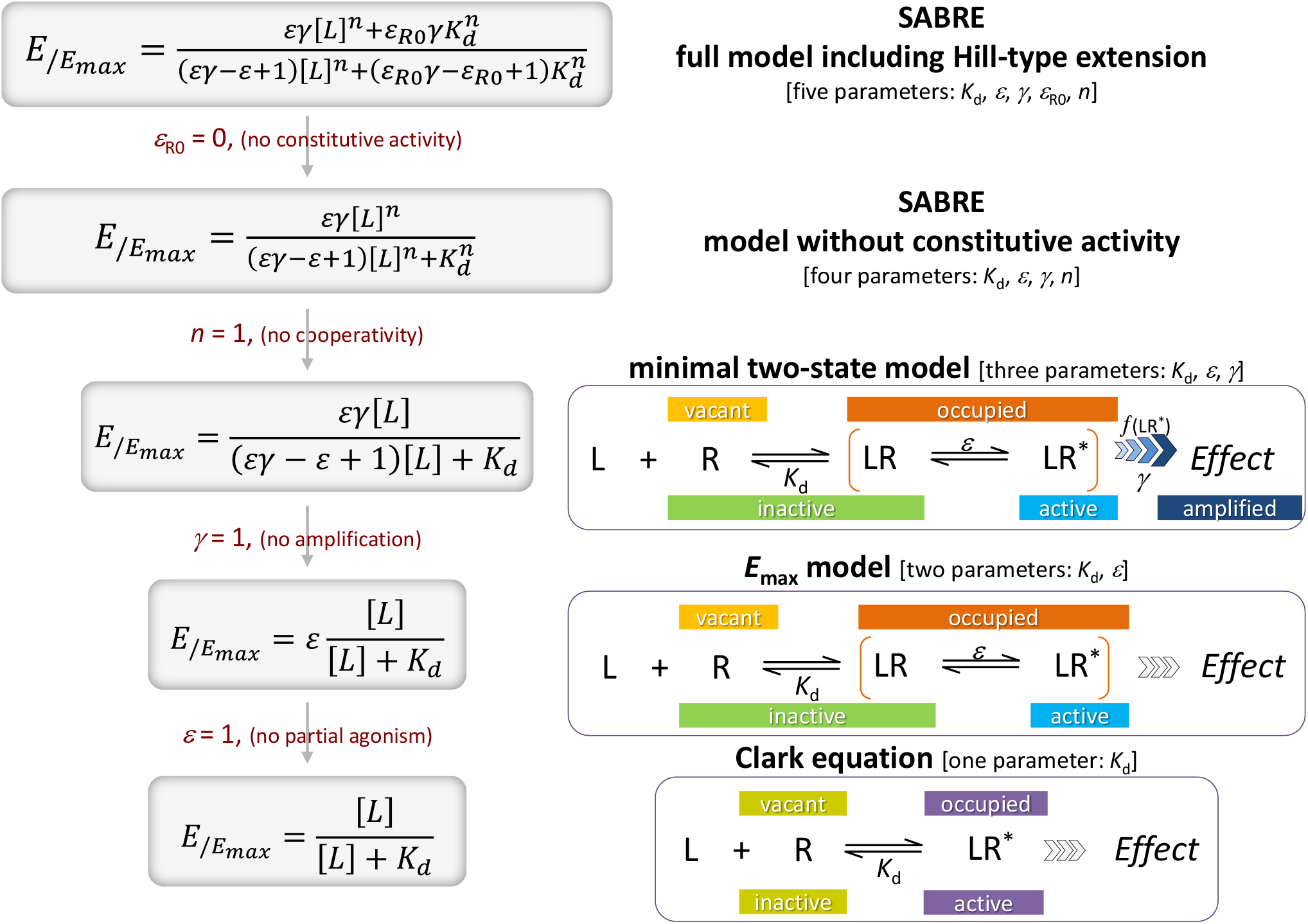
The relationship between ligand concentration [L] and receptor response (e.g., fractional response *f*_resp_ = *E*/*E*_max_) can be quite complex as illustrated here using the SABRE receptor model [1] starting with its most general form with Hill-type extension (top) and its consecutively nested simplifications down to the Clark equation (bottom) that can be obtained via consecutive restrictions of each parameter. Depending on the data, the simplest form (i.e., lowest number of adjustable parameters) should be used that can fit the data. Note that it is not necessary to impose *n* = 1 at the stage shown here, e.g., one can use a model with *K*_d_ and *n* as its adjustable parameters (i.e., a Hill instead of a Clark equation).

Thus, within the framework of SABRE, if there is no constitutive activity, half-maximal response (EC_50_) is observed at

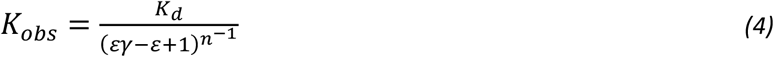

highlighting that, indeed, *K*_obs_ depends not just on receptor binding (i.e., occupancy determined by *K*_d_), but also on ligand efficacy (*ε*) and signal amplification (*γ*), not to mention cooperativity (*n*). It has also been shown that within the framework of SABRE (assuming *ε*_R0_ = 0 and *n* = 1), fractional response and occupancy are connected via the following function

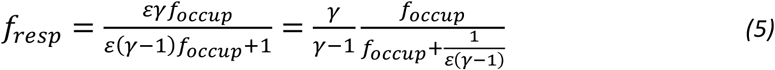

which can even be used for direct fitting if both (fractional) response and occupancy data are available (see [1]).

Considering these, methods that allow quantification of receptor binding (occupancy) from measurements of response (effect) data alone are of interest, as they can be used to characterize binding properties (e.g., *K*_d_) without having to perform explicit ligand binding experiments that typically require very different setups than response assays (e.g., use of labeled ligands or displacement of known labeled ligands). One so far underutilized possibility to achieve binding quantification this way is to acquire multiple concentration-response curves at different receptor levels, which can be obtained, for example, after different partial irreversible receptor inactivation (i.e., classic Furchgott method) [4, 5] or, more recently, by using different expression levels, and then fitting these response curves to obtain *K*_d_ estimates. The method of irreversible receptor inactivation introduced by Furchgott [4, 5] is particularly useful exactly because it allows the simultaneous estimation of affinity and efficacy when concentration-response functions can be established from the same preparation before and after partial irreversible receptor inactivation. It has been used since the late 1960s, albeit in a relatively limited number of works (e.g., ∼50 in PubMed). It was applied in several cases with various ligand series typically for G-protein coupled receptors (GPCRs), such as muscarinic [5-8], opioid [9-11], dopamine [12], 5-hydroxytryptamine (5-HT) [13, 14], and A_1_-adenosine [15, 16] receptors. An interesting variation that is now possible is to obtain responses not after different levels of partial inactivation, but with cells having different levels of receptor expression. A recent example of such data has been obtained with M_2_ and M_4_ muscarinic receptors and three ligands (carbachol, oxotremorine, and pilocarpine) at five different levels of expression in CHO cells [17].

Some type of fitting procedure is then needed to obtain the *K*_d_ estimates. Here, two alternative fitting methods are described to achieve this. One is a simple method via straightforward fitting of each response with sigmoid functions and estimation of *K*_d_ from the obtained *E*_max_ and EC_50_ values for each pair of control versus treated comparison. This is less error-prone than the original method of double-reciprocal fit proposed by Furchgott for his “null method” (see eq. 7) [4, 5] and is simpler than alternatives that require concentration interpolations, such as those proposed by Parker and Waud (see eq. 9) [18]. As it is very straightforward to implement using nonlinear regression software, which is now widely available, it is hoped that it will allow more widespread use of this so far underutilized approach to estimate binding properties (i.e., *K*_d_) or relative efficacies. The other is a more complex method that uses the above-mentioned SABRE model [1] to obtain a unified fit of the multiple concentration-response curves with a single set of parameters, one of which is *K*_d_. If sufficient data are available to allow reliable estimate of all parameters, SABRE can allow estimates obtained from all data points, which could be particularly advantageous if multiple concentration-response curves are available. For illustration, fit of simulated and experimental data are included with both models including for complex cases of multiple compounds of differing efficacies assessed at multiple receptor levels.

## METHODS AND MODELS

### Implementation and data fitting

All data fittings were performed in GraphPad Prism (GraphPad, La Jolla, CA, USA, RRID:SCR_002798) including those with the SABRE model as described before [1, 3]. Data used for fitting were normalized and considered as having no baseline (i.e., in the 0–100% range); parameters constrained to specific values are indicated for each fitting separately. Simulated data were generated with the same model in Prism using the “Simulate XY data” algorithm with 5% random error). Experimental data used for illustrations of model fit in Figure 3 were reproduced from reference [18]; those used for Figure 5 were obtained from Ehlert [19].

### Models for data fitting

#### Fit using double reciprocals and linear regression (Furchgott’s “null method”)

Following Furchgott, this is a method relying on comparing equal responses before and after treatment based on the hypothesis that equal stimuli should produce equal responses irrespective of the actual shape of the transduction function linking the stimulus *S* to the generated response (effect) *E*. If one assumes that inactivation reduces the number of total receptors to a *q* fraction of the original, [R_tot_]’=*q*[R_tot_], while response continues to be generated via the same transduction function, the concentrations [L] and [L]’ that produce the same effect pre- and post-inhibition must create the same *stimulus S* (using the original terminology of Furchgott). Hence, if *S* is proportional with the concentration of occupied receptors [LR], *S* = *ϵ*_F_[LR], and the concentration of occupied receptors [LR] is connected to that of the ligand [L] via the standard hyperbolic function (law of mass action), equal pre- and post-inactivation responses imply that

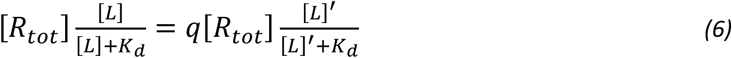

This can be rearranged into the linear relationship between the reciprocals of equiactive concentrations that is the basis of the original Furchgott method introduced in the late 1960s [4, 5, 20]:

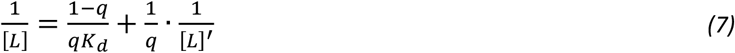

The slope and intercept of this line allow the simultaneous determination of *q* (fraction of receptor inactivated) and *K*_d_ (receptor affinity). It was considered an important advantage that this Furchgott analysis is a null method approach that makes no assumption regarding the nature of the transduction function other than it stays the same after partial inactivation of R_tot_. It has, however, several implicit assumptions [5] including that response is produced via only one transduction system (i.e., no poly-pharmacology) and that receptor occupation follows a classic law of mass action (i.e., hyperbolic function as shown in eq. 2 with no cooperativity, Hill slope *n* = 1). One of its important drawbacks is that it relies on linear regression using a double-reciprocal plot, and, hence, just as the Lineweaver-Burke plot, it is prone to errors typical for such plots: it tends to amplify the errors of the measurements at low concentrations (where 1/[L] is large), as those points will weigh more heavily in the corresponding linear regression, so that the very worst data are emphasized the most, as the smallest measured numbers have the largest influence on the regression due to the use of reciprocal values [21]. Another important drawback is that, except for rare lucky coincidences, actual measurements at equiactive concentrations are not available, so that some interpolation method has to be used to estimate at least one of the concentration values producing the same response.

#### Fit using hyperbolic functions and interpolated concentrations (Parker-Waud)

Following the suggestion of Parker and Waud in the early 1970s, an often used approach to overcome the problems of this double-reciprocal method was to (1) fit the control (untreated) data with a standard hyperbolic function, (2) use this to obtain the needed interpolated [L] concentrations, (3) fit those using another nonlinear regression for the equation connecting [L] and [L]’, and then (4) use the parameters of this fitting to estimate *q* and *K*_d_ [18]. Specifically, first a nonlinear regression is used to fit the pre-inactivation control data with a classic sigmoid (Hill) function similar to eq. 1

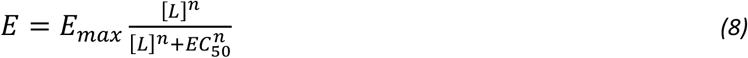

Next, this is used to interpolate the [L] concentrations that would produce the same responses pre-inactivation as those measured post-inactivation at the [L]’ values tested, and then another nonlinear regression is used to fit the corresponding [L] and [L’] pairs to the hyperbolic equation connecting them from which *q* and *K*_d_ can be obtained (essentially, the reverse of eq. 7):

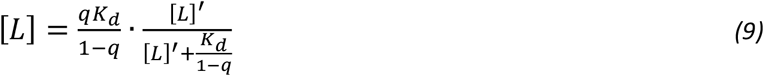

These make fitting a cumbersome and multi-step process – quite possibly a main reason behind the relatively rare use of this otherwise very informative approach to estimate *K*_d_ from response data alone. A nested version of this, where eq. 9 is used to replace [L] in eq. 8 and fit *E*’ as a function of L’ in parallel with fitting *E* to [L] using eq. 8, was proposed by James and co-workers in the late 1980s [22].

#### Fit using individual sigmoid functions (present simple method)

To overcome these cumbersome fittings, here, first a simple alternative is proposed with fitting that can be easily done with software tools for nonlinear regression that are now widely available and used to directly obtain estimates for *K*_d_ and *q*. One possible alternative to avoid the use of double-reciprocal regression and the need for concentration interpolations is to simply fit both sets of activity data with standard hyperbolic curves (i.e., sigmoid on semi-log scale) and use the parameters from this to obtain *q* and *K*_d_. This was avoided in the original approach to not make any assumptions on the shape of the transducer function. However, hyperbolic functions are good representations of typical concentration-response functions, and because of the extrapolations needed to obtain concentrations producing equipotent activities, fit in practice typically involved the assumption that at least one of the responses had some defined (typically, hyperbolic) functional form (sigmoid on log-scale). Hence, more reliable parameters can be obtained by using nonlinear regression to directly fit both responses with sigmoid curves and use those to estimate *q* and *K*_d_ values. Assuming straightforward hyperbolic functions (i.e., *n* = 1) both before and after inactivation

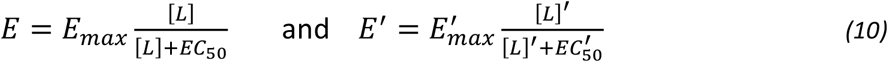

the ligand concentrations [L] and [L]’ causing effects *E* and *E*’ are obtained as:

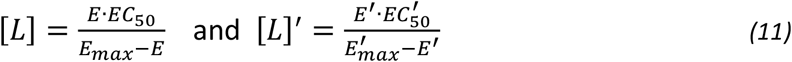

This can be used to link the reverse of concentration 1/[L] to that of the reverse of equiactive equivalent 1/[L]’ by replacing *E* with *E*’, which are equal here:

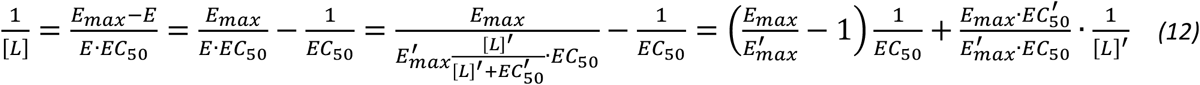

By comparing the slope and intercept of this with those of the double-reciprocal linear relationship that is the basis of the Furchgott method (eq. 7), one can see that

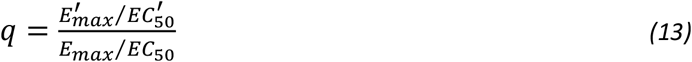

and (after some transformations using this):

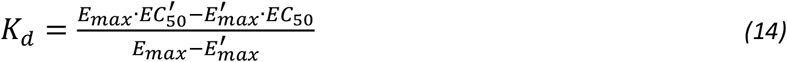

Hence, this method of fitting two sigmoid response functions can provide faster and more reliable estimates for *q* and *K*_d_ than the previous more cumbersome methods requiring interpolated concentrations. If multiple compounds were assayed in the same system, relative efficacies can also be estimated once *K*_d_ estimates are available by comparing fractional occupancies *f*_occup_ (calculated from these *K*_d_ values) that cause the same (fractional) effect [5]. A simple formula for estimating relative efficacies using the *E*_max_, EC_50_, and *K*_d_ values obtained via the sigmoid fitting is (see derivation assuming sigmoid response and occupancy functions in Supporting Information, Appendix 1 and with SABRE’s formalism below using eq. 26):

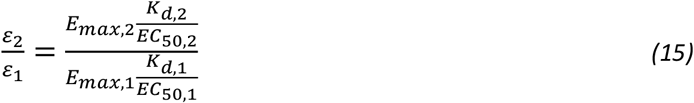

While this method of fitting sigmoid functions appears to abandon the apparent elegance of the “null method” of making no assumptions regarding the actual shape of the transduction function, in practice, it makes little difference as sigmoid (or some other similar) functions had to be used anyway for the needed data interpolations. Note also that *q* as estimated with this method is the same as the ratio of equiactive molar ratios (EAMR; i.e., ratio of molar concentrations that produce the same response at low enough concentrations), later termed intrinsic relative activity (*RA*_i_) introduced by Ehlert and co-workers, which is the ratio of *E*_max_/EC_50_ values for a Hill slope of 1 [23, 24]:

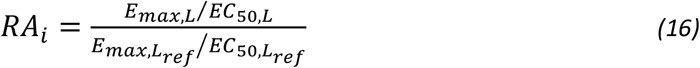

This became the basis of bias quantification methods that commonly rely on calculating the shift ΔΔlog(*E*_max_/EC_50_), which is estimated as ΔΔlog(τ/*K*_D_) if the operational model is used, versus a selected reference compound, e.g., a logarithmic bias factor is obtained as (log(*E*_max,P₁,L_/EC_50,P₁,L_)– log(*E*_max,P₂,L_/EC_50,P₂,L_)) – ((log(*E*_max,P₁,Lref_/EC_50,P₁,Lref_)–log(*E*_max,P₂,Lref_/EC_50,P₂,Lref_)) [25-28]. In fact, using *E*_max_·*K*_d_/EC_50_ from eq. 15 derived here as a measure to compare relative activities or efficacies makes more sense than using *E*_max_/EC_50_ from eq. 16, which are being used as *intrinsic relative activities* (*RA*_i_ or EAMD), as the former is a dimensionless quantity (%), while the latter is not (mol^−1^). When used for the same compound (e.g., comparing activities at different receptor levels, i.e., Furchgott method, or for different diverging downstream pathways, i.e., biased agonism), the comparison in eq. 15 reduces to the same as eq. 16, as *K*_d_s are the same and are eliminated from the ratio.

#### Fit using SABRE (present, unified method)

Finally, a more complex approach is also described that uses the recently introduced multiparametric SABRE receptor model to achieve a single unified fit of the multiple concentration-response curves and, hence, a *K*_d_ estimate based on all data available. The SABRE quantitative receptor model in its most general form uses a total of five parameters: three characterizing the interaction of the ligand with the receptor (binding affinity, *K*_d_, efficacy, *ε*, and Hill coefficient, *n*) and two characterizing the receptor signaling pathway (amplification, *γ*, and constitutive activity, *ε*_R0_), (Figure 1) [1-3]:

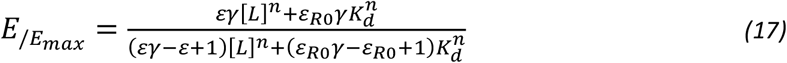

With the incorporation of separate parameters for signal amplification and (intrinsic) efficacy, SABRE can fit concentration-response functions even for complex connections between fractional response (*f*_resp_) and occupancy (*f*_occup_) [1]. An important advantage compared to other complex receptor models, such as those based on the operational (Black and Leff) model [29, 30], is that SABRE’s equations can be collapsed into consecutive simplified forms by fixing the parameters as special values (Figure 1), and these can and should be used on their own when adequate [1]. Furthermore, SABRE can accommodate experimental *K*_d_ data and, hence, connect experimentally determined fractional occupancy and fractional response data, which is not possible within the framework of the original operational model. Note also that the operational model is not well suited to fit Furchgott type data; it could not provide independent estimates of agonist affinity and efficacy [31].

As discussed before [1], the simplest form of SABRE that still provides adequate fit should be used to avoid over-parametrization, and selection of the model to be used should also be guided by the consideration that reliable fitting and well-defined parameter values can only be achieved if there are at least 5–10 (well-distributed) data points for each adjustable parameter [32-34]. As data here do not involve constitutive activity, the general form of SABRE that assumes no constitutive activity (*ε*_R0_ = 0) but still allows a Hill-type extension was used for all fittings (Figure 1, second from top; also eq. 3):

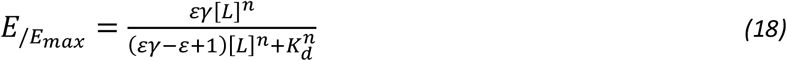

As shown before, SABRE can be used for the quantitative modeling of responses measured after partial irreversible receptor inactivation [3]. The loss of total receptors due to irreversible inhibition, [R_tot_]’=*q*[R_tot_], leads to a corresponding loss in the concentration of active receptors [LR^*^] responsible for generating the response and, hence, a loss in response. Assuming that the inactivation does not affect the post-receptor signal amplification function, the fractional response after inactivation in the formalism of SABRE becomes:

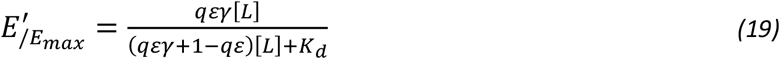

Thus, a *q*-fold decrease in R_tot_ translates into an apparent *q*-fold reduction of efficacy as used in SABRE: *ε*’=*qε*. Two illustrations for the use of SABRE with this formalism of “fractional efficacy” *ε*’ and unified amplification (*γ*) and *K*_d_s to fit data from Furchgott type experiments have been provided in [3]: one with a series of compounds acting at the dopamine receptor, which was inactivated by the irreversible antagonist *N*-ethoxycarbonyl-2-ethoxy-1,2-dihydroquinoline (EEDQ) [12], and one with the muscarinic agonists carbachol and oxotremorine and receptor inactivation with phenoxybenzamine (PHB) [35]. Further examples with simulated and experimental data are presented below including with muscarinic agonist activities measured in rabbit myocardium after different levels of inactivation caused by an irreversible muscarinic antagonist (BCM).

The framework provided by SABRE can also be used to confirm equations 13–15 obtained for the fitting of individual sigmoid functions above. Using the rearranged form of SABRE (shown in eq. 3)

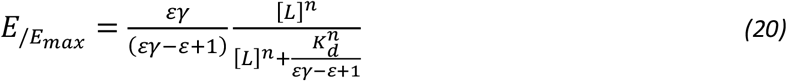

it is clear that half-maximal response (EC_50_, *K*_obs_) is observed at

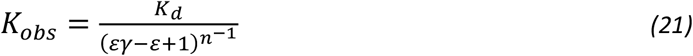

and the maximum fractional effect is

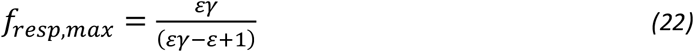

Thus, the term of interest for ratio as intrinsic relative activity (equiactive molar ratio; eq. 16), which also corresponds to the “transduction coefficient” τ/*K*_D_ of the operational model, is *εγ*/*K*_d_ in SABRE assuming *n* = 1 (see Appendix 2 of [1]):

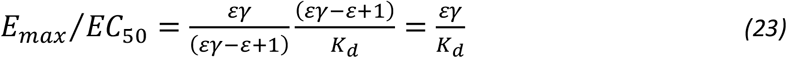

Thus, fitting of two individual sigmoid curves and using eq. 13 indeed reproduces *q* in the formalism of SABRE too with the modified efficacy, *ε*’=*qε*:

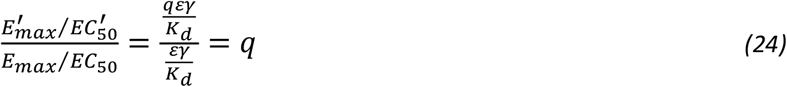

Similarly, eq. 14 reproduces *K*_d_:

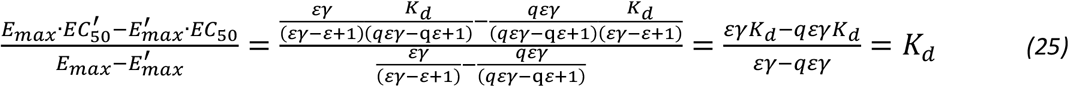

If multiple compounds are assayed in the same system, eq. 23 can be used to obtain an estimate of relative efficacies for two compounds (*ε*_2_/*ε*_1_) even with the sigmoid fitting approach once *K*_d_s have been calculated from eq. 14. Since pathways and hence *γ*s are the same, the ratio of two efficacies (i.e., relative efficacy versus a reference compound) as expressed from eq. 23

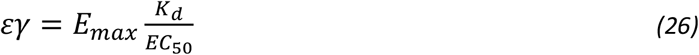

reproduces exactly the one shown in eq. 15 earlier.

## RESULTS AND DISCUSSION

### Illustration 0: Fit of response data with sigmoid function and SABRE

First, fitting of response only data including partial agonists is shown to illustrate the parallel between use of classic sigmoid functions (i.e., eq. 1) and that of SABRE (eq. 18) with its simplified form that also assumes no amplification (*γ* = 1), hence, has only three adjustable parameters, *K*_d_, *ε*, and *n*:

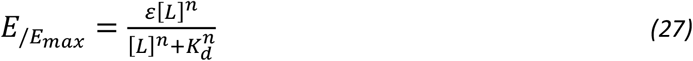

Simulated data generated for three different hypothetical agonists (with a 5% random error) was used as shown in Figure 2. Fitting with SABRE produced the very same parameters as fitting with classic sigmoid functions (i.e., “log(agonist) vs. response - Variable slope (four parameters)” in GraphPad Prism) with the same constrains (Bottom = 0, HillSlope shared values for all datasets) and nicely reproduced the log *K*_d_ and maximum fractional response *f*_resp,max_ values used to generate the data: −7.998_±0.017_, −6.211_±0.023_, and −7.564_±0.056_ for the log *K*_d_ values used of −8.000, −6.200, and −7.500; and 0.989_±0.007_, 0.765_±0.008_, 0.308_±0.007_ for the maximum responses used of 100%, 75%, and 30% (Supplementary information, Table S1). For further generalizability, data corresponding to a Hill slope of *n* = 2 were used here, and fitting was done by constraining *n* to the same value for all compounds resulting in a fitted value of 1.828_±0.090_.

**Figure 2.**
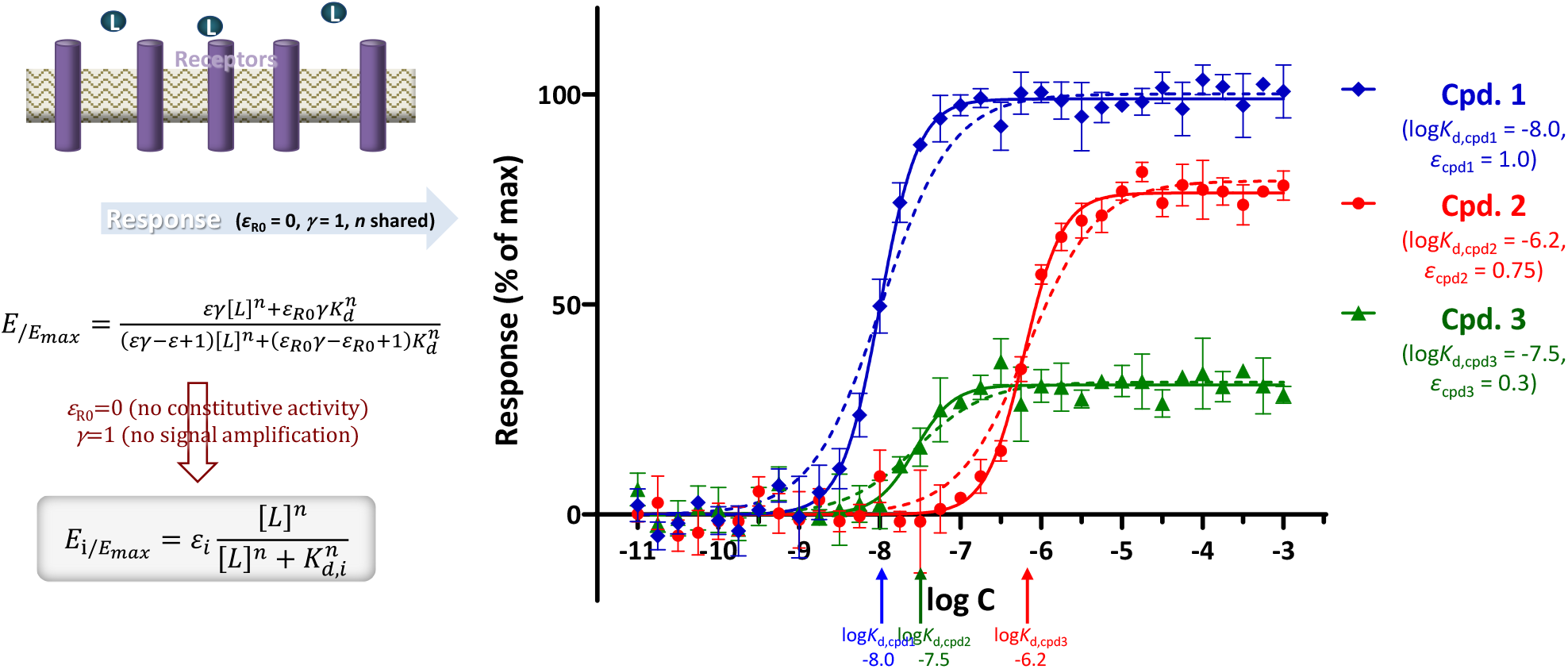
Fit of response only data for full and partial agonists with SABRE using its form shown in eq. 20 that allows non-unity Hill slope. Simulated data (symbols) for three different compounds were generated in GraphPad Prism as described in the Methods (“Simulate XY data” algorithm with 5% random error) and then fitted with SABRE (lines) using the following constrains: *ε*_R0_ = 0, *γ* = 1, *n* shared. Fit with classic sigmoid functions, “log(agonist) vs. response - Variable slope (four parameters)” with Bottom = 0 and HillSlope shared as constrains, produces the exact same fit (Supplementary information, Table S1) with overlapping lines. Dashed lines show fit if only unity Hill slopes are allowed; data here correspond to a Hill slope of *n* = 2 (note the wide range of data included that covers nine orders of magnitude).

Hence, if adequately constrained (*ε*_R0_ = 0, *γ* = 1), SABRE can perform the same role as the widely used Hill or Clark (*n* = 1) equation and can be used to derive straightforward EC_50_ (*K*_obs_) values. However, it can do much more than that if its other parameters are released and can be adequately fitted. For example, it can fit the same type of response data, but with integration of occupancy (*K*_d_) data obtained from a different, independent assay in the same system. In other words, it can connect fractional response, *f*_resp_, and fractional occupancy, *f*_occup_, data even in complex cases where the fractional response can be either ahead or behind the fractional receptor occupancy. Fit with SABRE can be considered consistent if adequate fit can be obtained with well-defined log *K*_d_ values (that are consistent with their experimental values if those are available) and a single amplification parameter *γ* characterizing the system (pathway) as a whole. Examples for this have been shown before for simulated [1] as well as experimental data including among others imidazoline type *α*-adrenoceptor agonists data [36] often used as textbook illustration of mismatch between *f*_resp_ and *f*_occup_ [3, 37]. Contrary to SABRE, the classic two-parameter (τ, *K*_D_) operational model [29, 30] can neither be simplified back to the Hill or Clark equation by constraining its τ parameter nor used to connect the response to independently determined binding data (*K*_d_) [1, 3]. A “special edition” extension of the operational model “with given *K*_d_ values” introduced by Rajagopal and coworkers for bias quantification can use experimental *K*_d_ values as its *K*_D_s (instead of fitted *K*_D_s) to constrain the regression [25, 27], but this required introduction of an additional scaling factor *α* making it a three- and not a two-parameter model (see [1] for details and correspondence between the parameters of these models).

### Illustration 1: Guinea pig ileum treated with dibenamine

A first illustration of fit of receptor inactivation data (Figure 3) is provided with the relatively simple experimental data obtained with guinea pig ileum and used by Parker and Waud to introduce their fitting method for concentration-responses following partial irreversible receptor inactivation [18]. Because of the limited number of data points available, fits were done with the Hill slopes fixed as unity (*n* = 1), especially since even if released, the fitted values for *n* were not significantly different from 1. The two fits here (individual sigmoid curves and SABRE) produced very similar results, i.e., completely overlapping curves in Figure 3 and predicted fraction of receptor inactivated *q* = 0.40 and receptor affinity log *K*_d_ = −4.11 for both, which are in good agreement with the original estimates obtained by Parker and Waud (Supplementary information, Table S2).

**Figure 3.**
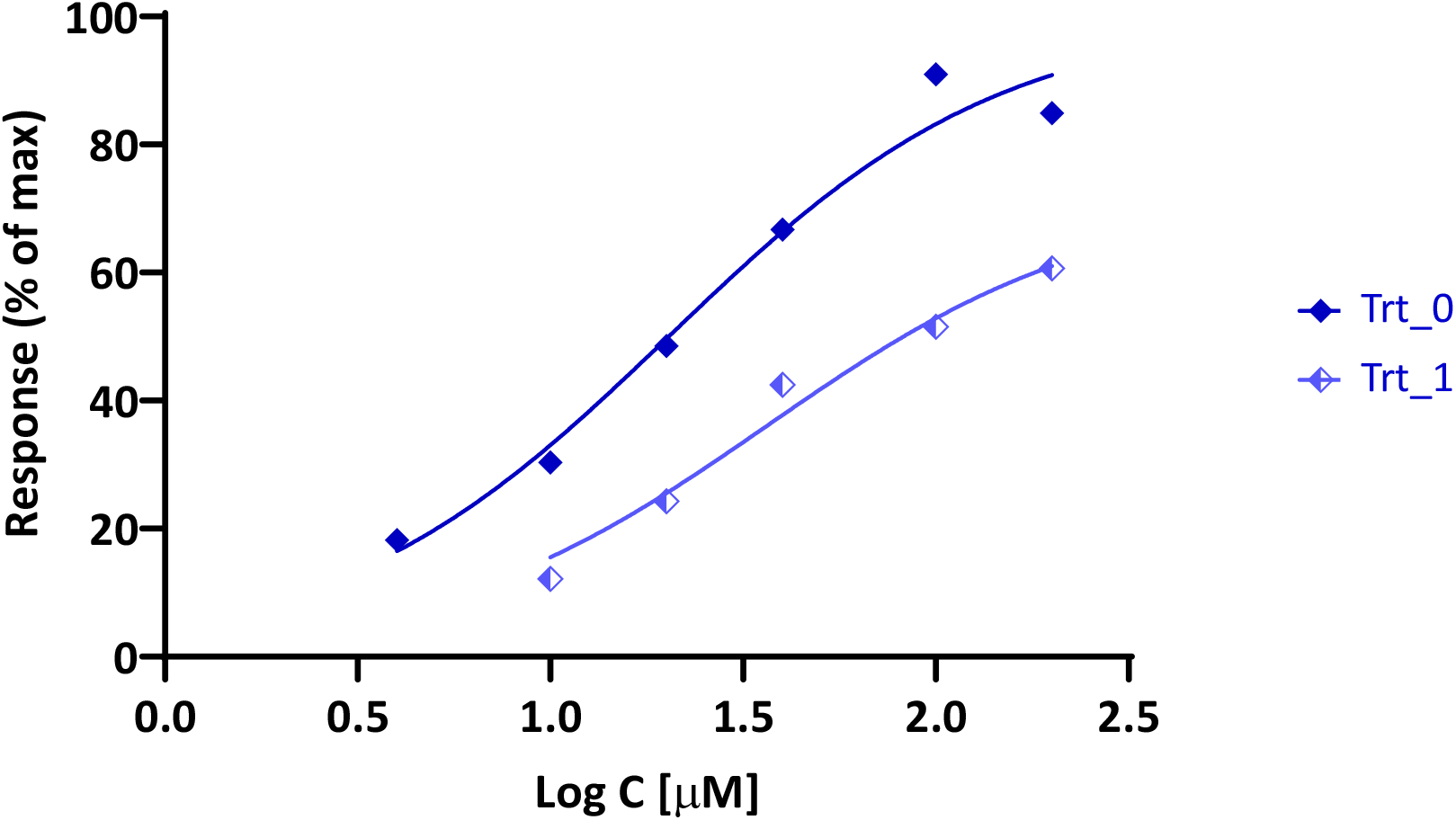
Fit of concentration-response curves from guinea-pig ileum preparations treated with heptyl(trimethyl)azanium, *n*-C_7_H_15_N^+^(CH_3_)_3_, before (Trt_0) and after (Trt_1) administration of the irreversible antagonists dibenamine (data after [18], normalized). Fit with SABRE and with individual sigmoids overlap and do not separate on the graph.

### Illustration 2: Complex simulated receptor inactivation data

An example with simulated data generated to illustrate that full agonists can become partial ones following receptor inactivation and even the order of apparent potencies can change is shown in Figure 4 with parameters summarized in Supplementary information, Table S3. The three different responses (corresponding to no treatment, Trt_0, and two inactivation treatments, Trt_1 and Trt_2) obtained for each of the three compounds (color coded as blue, red, and green for compounds 1, 2, and 3, respectively) are shown as a function of log concentration individually and in combination in Figure 4. Unified fit with SABRE nicely reproduced the amplification (*γ* = 2,000), receptor inactivation (*q*_1_ = 0.030, *q*_2_ = 0.001), binding affinity (log *K*_d_: −5.0, −7.0, and −5.0), and ligand efficacy (*ε*: 1.00, 0.02, and 0.30) values used to generate the data (Table S3B). For example, the estimates obtained from fitting were - −5.031_±0.043_, −6.949_±0.031_, and −4.955_±0.044_ for the log *K*_d_ values used to generate the data of −5.000, −7.000, and −5.000. Fit with individual sigmoids also produced acceptable estimates (one for each inactivation treatment, which can be averaged for overall estimates; Table S3A). Note, however, that on such multiple inactivation data, SABRE provides more reliable estimates overall because of the unified fit of all data, if it can provide one. Here, individual fit with sigmoids could not obtain sufficiently reliable estimates for *K*_d,Cpd1_ from treatment 1 data (Cpd1_Ttr_1_) as the two maxima were very similar, i.e., log *K*_d_ estimate of −4.589 vs the actual −5.000, and for *K*_d,Cpd2_ from treatment 2 data (Cpd2_Trt_2_) where the responses were too small for reliable fitting, i.e., log *K*_d_ estimate of −7.578 vs the actual −7.000. These values that correspond to *K*_d_ values that are 3–4-fold off, as well as the *q*_2_ value derived from the data of Cpd2 that is also about 4-fold off, are shown in italic in Table S3A. For the same reason, the relative efficacy estimates calculated using eq. 15 are also almost two-fold off compared to the values used to generate the simulated data, unless they are calculated omitting these values (Table S3A; italic vs non-italic values in row denoted with asterisk).

**Figure 4.**
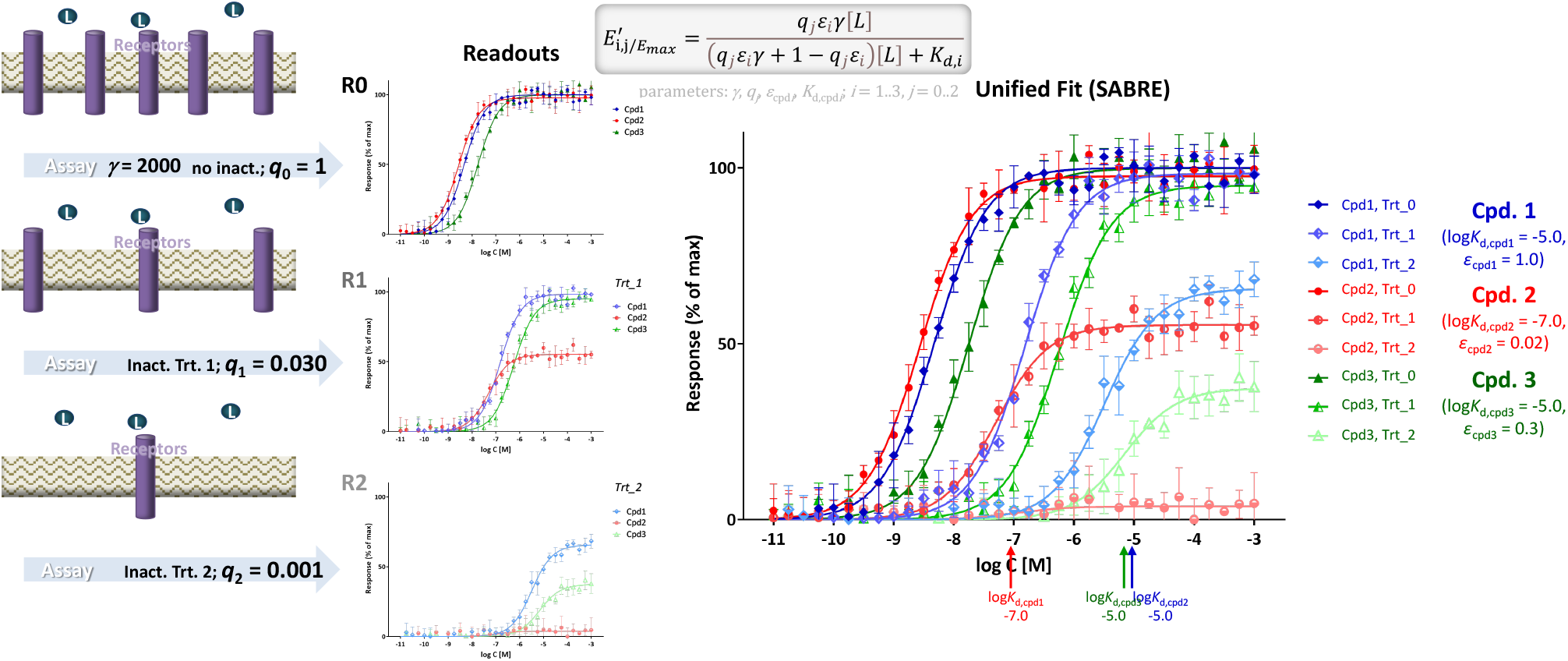
Fit of responses measured after partial irreversible receptor inactivation (Furchgott method) with SABRE using unified pathway amplification (*γ*) and ligand affinity (*K*_d_) parameters. Simulated data (symbols) for three different compounds (Cpd. 1, 2, and 3) and two different inactivation levels (Trt_1 and Trt_2) were generated in Prism as described in Methods with the parameter values as shown (three compounds with corresponding log *K*_d,i_s and *ε*_*i*_s, *i* = 1, 2, and 3 at three different inactivation levels with corresponding *q*_*j*_s, *j* = 0, 1, and 2; 5% random error) and fitted with SABRE (lines) using a total of nine parameters (Table S3B). Data were selected to illustrate that receptor inactivation can change full agonists into partial ones and even the order of apparent potencies – compare compound 2 (red) and 1 (blue) at Trt_0 vs Trt_2.

### Illustration 3: Muscarinic activity in rabbit myocardium

A further detailed illustration with experimental data is included using the muscarinic receptor-mediated inhibition of adenylate cyclase activity measured in rabbit hearts (Figure 5). Activity of three muscarinic agonists, namely oxotremorine-M, oxotremorine, and BM5 (N-methyl-N-(1-methyl-4-pyrrolidino-2-butynyl)acetamide), was determined in perfused rabbit myocardium homogenates in the absence and presence of the irreversible muscarinic antagonist benzilylcholine mustard (BCM; 0, 1.0, and 10.0 nM, 15 min) [19]. Unified fit with SABRE of 81 data points from all 9 response data sets (3 compounds assessed at 9 concentrations each at 3 different pretreatments; 3×9×3 = 81) using only 10 parameters (3×*K*_d_, 3×*ε, γ, n, q*_1_, *q*_2_) resulted in good overall fit (Figure 5) accounting for 99.3% of the variability in the data (*r*^2^ = 0.993) with an amplification of *γ* = 7.3_±1.6_ (Table S4B). Here, individual fit with sigmoids (Table S4A) also produces good fit, in fact, slightly better one (*r*^2^ = 0.994; SSE 474.1 vs 550.0 with SABRE); however, it requires a total of 19 parameters (2, *K*_d_ and *E*_max_, for each of the 3 compounds in 3 treatments plus *n*; i.e., 2×3×3+1 = 19). According to the fit with SABRE, BCM caused fractional inhibitions (*q*) of 0.406 and 0.065 for the 1.0 and 10.0 nM pre-treatments, respectively (to be compared with estimates of 0.380 and 0.020 obtained in the original publication using analysis of equiactive agonist concentrations [19]). Fit with sigmoids produced averaged *q* estimates of 0.374 and 0.067, similar to those from SABRE; however, the three different estimates from the three compounds for *q*_2_ were a bit different covering an approximate two-fold range from 0.051 to 0.092 (Table S4A). For the three compounds, log *K*_d_ estimates from SABRE came out as −5.27, −5.85, and −6.31 for oxotremorine-M, oxotremorine, and BM5, respectively (corresponding to *K*_d_s of 5.3, 1.4, and 0.5 *µ*M) in nice agreement with the values measured by competition assays (4.0, 1.1, and 0.2 *µ*M) as well as those estimated from fitting (7.5, 2.2, and 0.2 *µ*M) in the original work [19]. Efficacy (*ε*) estimates from SABRE were 1.0, 0.75, and 0.13 – in reasonable agreement with the original estimates of 1.0, 0.52, and 0.04 obtained by Ehlert using equiactive concentrations [19]. Relative efficacies estimated using the sigmoid approach for fitting and then eq. 15 were also similar to those from SABRE being 0.64 for oxotremorine and 0.12 for BM5 compared to oxotremorine-M as reference (Supplementary information, Table S4A).

**Figure 5.**
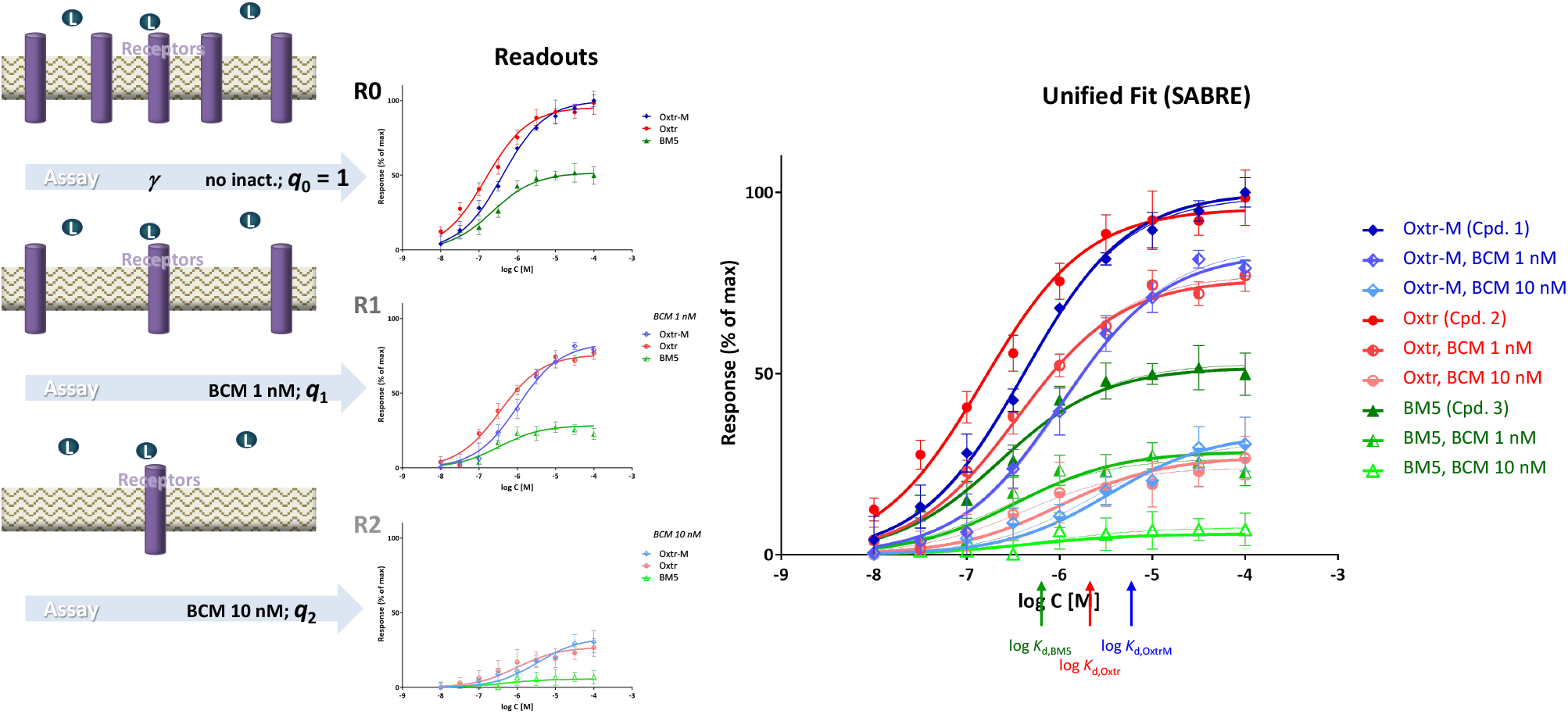
Fit of experimental responses obtained for three muscarinic agonists (oxotremorine-M, oxotremorine, and BM5) measured before and after partial irreversible receptor inactivation with BCM (0, 1.0, and 10.0 nM, 15 min) [19]. Unified fit of all 81 data points done with SABRE with 10 parameters (3×*K*_d_, 3×*ε, γ, n, q*_1_, *q*_2_; Table S4B) is shown in color-coded thick lines. For comparison, fit with individual sigmoids is also included (thin lines; 19 parameters: 3×3×*K*_d_, 3×3×*E*_max_, *n*; Table S4A).

## CONCLUSION

In conclusion, multiple response only data measured at different receptor levels can be used to quantify receptor binding without having to perform ligand binding / displacement experiments; however, careful fitting is required to separate binding affinity from efficacy (i.e., partial agonism) and signal amplification (“receptor reserve”). Here, two fitting methods were described that are both straightforward to use with widely available nonlinear regression software. One is a simple method that relies on fitting of individual sigmoid functions to provide *K*_d_ estimates from the obtained *E*_max_ and EC_50_ values. Since it requires no interpolation and needs only easily obtainable parameter estimates, e.g., *K*_d_ = (*E*_max_·EC’_50_ − *E*’_max_·EC_50_)/(*E*_max_ − *E*’_max_), it is hoped that it will allow more widespread use of this so far underutilized approach to estimate binding affinities. The other method uses the SABRE model to obtain a unified fit of the multiple concentration-response curves with a single set of parameters. This could be particularly advantageous if multiple concentration-response curves are available and a consistent fit can be obtained, as it provides binding affinity, *K*_d_, as well as efficacy, *ε*, and amplification *γ*, parameters derived based on the entire set of data.

## SUPPLEMENTARY INFORMATION

Supplementary information for this article includes Tables S1–S4 with detailed parameters from all fittings and Appendix 1 with a derivation of relative efficacies (eq. 15) assuming sigmoid response and occupancy functions.

## CONFLICT OF INTERESTS

The author declares that the research was conducted in the absence of any commercial or financial relationships that could be construed as a potential conflict of interest.

## AUTHOR CONTRIBUTIONS

PB is the sole author; he originated the project, performed the calculations and data fittings, and wrote the manuscript.

## FUNDING SOURCES

N/A

## ACKNOWLEDGMENT

The author would like to thank Fred Ehlert for providing the original experimental data obtained for the muscarinic agonists used for illustrations here and for his insightful comments and input on an early version of this manuscript.

## ABBREVIATIONS

BCM: benzilylcholine mustard
BM5: N-methyl-N-(1-methyl-4-pyrrolidino-2-butynyl)acetamide)
GPCR: G-protein coupled receptor
SABRE: present model (with parameters for Signal Amplification, Binding affinity, and Receptor activation Efficacy)
SSE: sum of squared errors.

## DATA AVAILABILITY STATEMENT

Data used for illustrations of model fit are either simulated data generated as described or reproduced from previous publications as indicated in the corresponding figures. The datasets generated and/or analyzed are available from the corresponding author upon reasonable requests.

## SUPPLEMENTARY INFORMATION

## Supplementary Tables

**Table S1.**
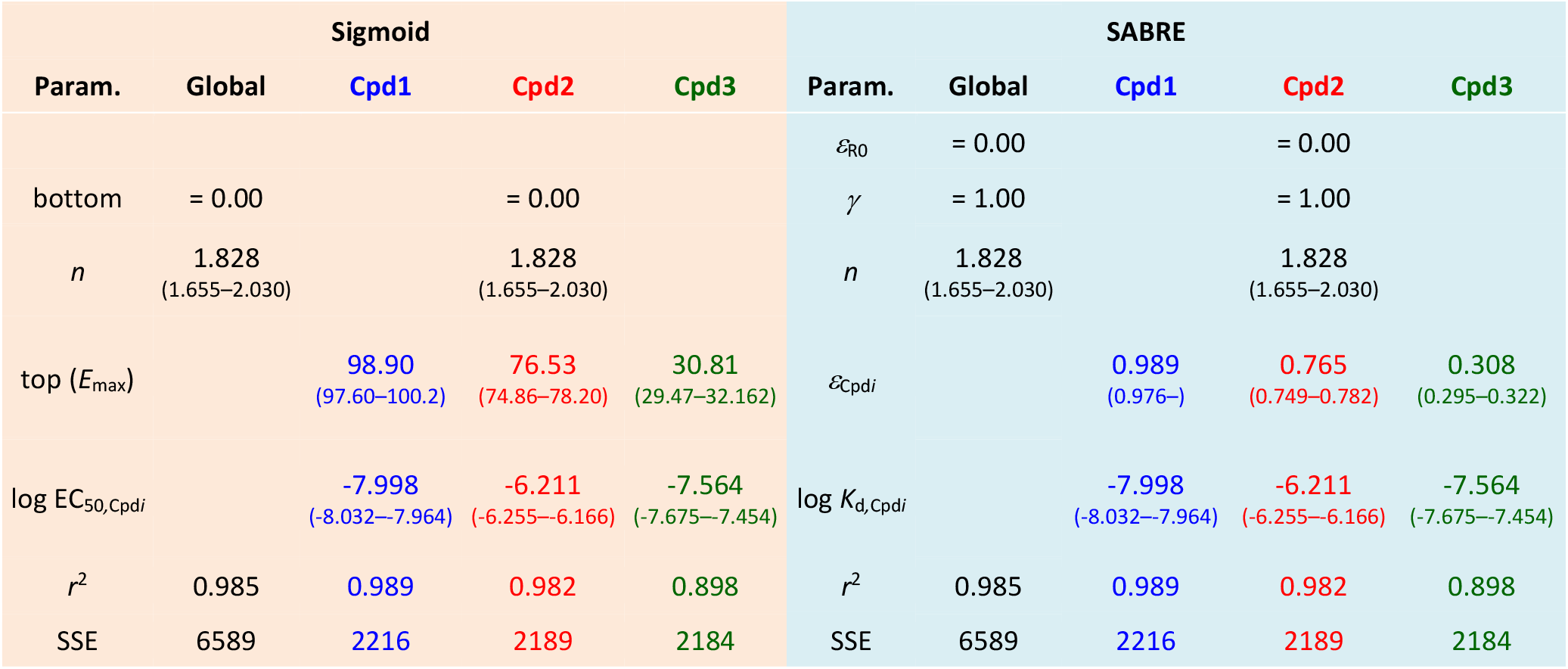
Parameters from fitting of data from Figure 2 with classic sigmoid model (log agonist vs response) and SABRE. Calculated parameters obtained using GraphPad Prism implementation for fitting are shown with their 95% confidence intervals (CI) and descriptors of the quality of fit (correlation coefficient, *r*^2^, and sum of squared errors, SSE). Fit with SABRE is identical to that obtained with the standard sigmoid Hill model, e.g., “log(agonist) vs. response - Variable slope (four parameters)” model in Prism (with shared *n* and bottom restricted to 0).

**Table S2.**
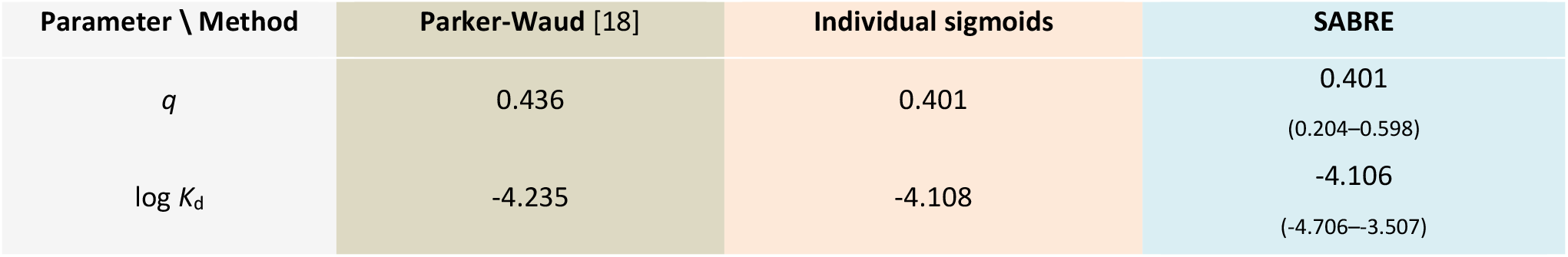
Fitting results for data from Figure 3 (guinea-pig ileum preparations treated with heptyl(trimethyl)azanium, *n*-C_7_H_15_N^+^(CH_3_)_3_ before and after inactivation by the irreversible antagonists dibenamine (data after [18]).

**Table S3.**
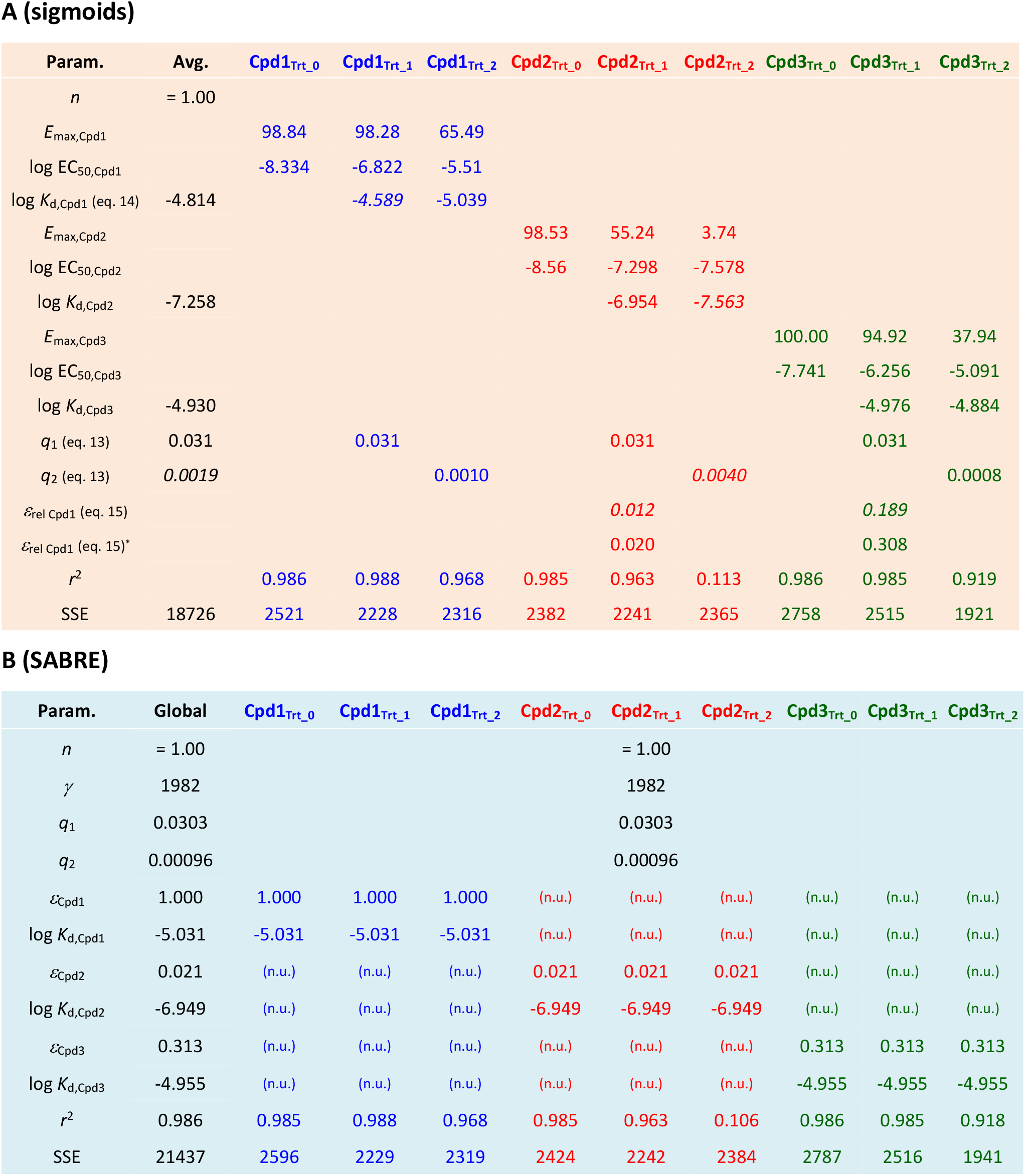
Parameters derived from fitting with individual sigmoid functions (**A**) and SABRE (**B**) of data from Figure 4. Calculated parameters obtained from fitting using GraphPad Prism are shown with descriptors of the quality of fit (correlation coefficient, *r*^2^, and sum of squared errors, SSE). For this fit, the Hill coefficient (slope) was fixed to unity as indicated (*n* = 1) in both cases (even if the sigmoid model is derived for *n* = 1 only). For SABRE in B, n.u. denotes parameters not used for the corresponding column.

**Table S4.**
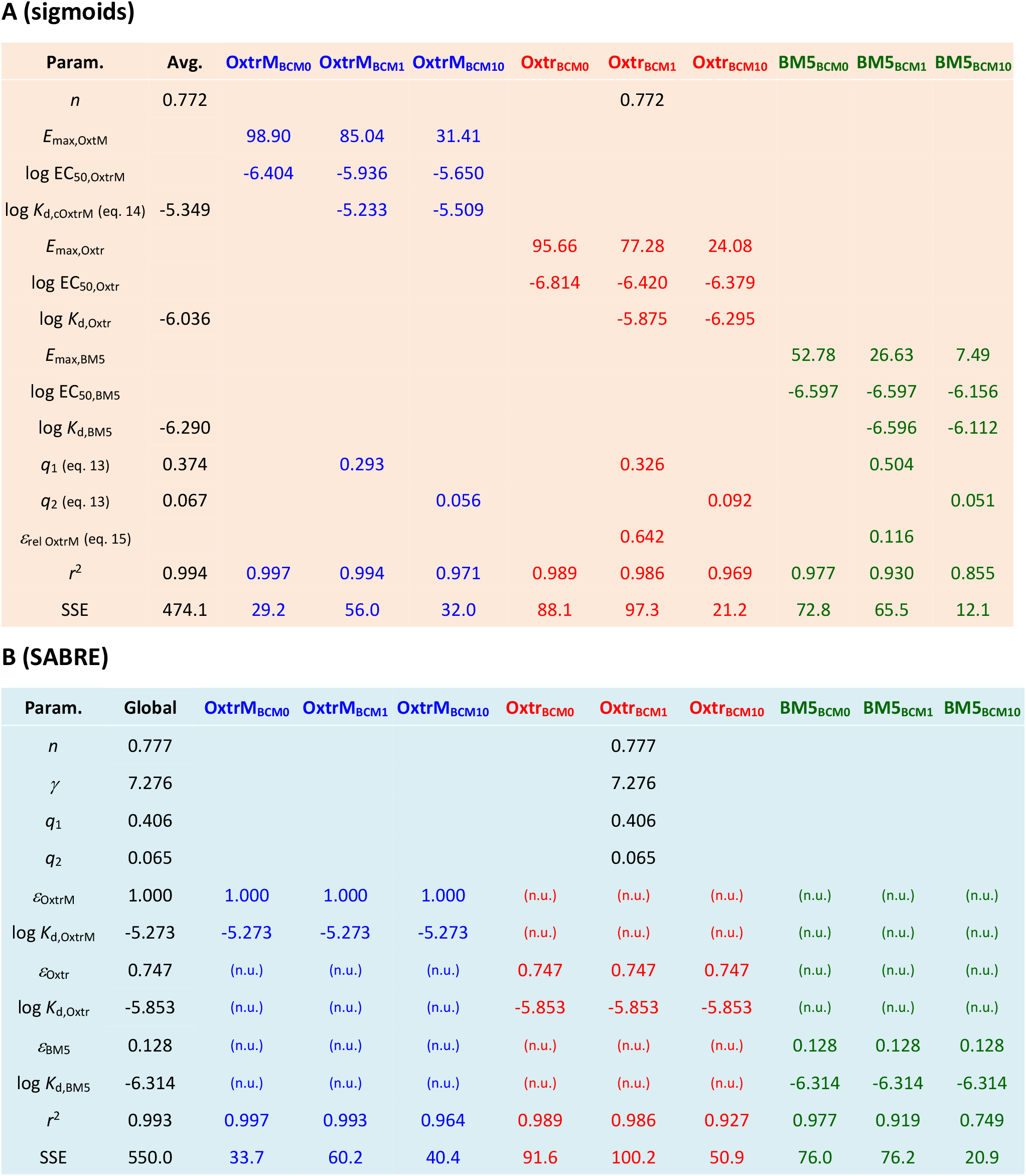
Parameters derived from fitting with individual sigmoid functions (**A**) and SABRE (**B**) of data from Figure 5 (muscarinic activity in rabbit myocardium). As before, calculated parameters obtained from fitting using GraphPad Prism are shown with descriptors of the quality of fit (*r*^2^ and SSE). For the sigmoid fitting, to obtain a better fit, which is also more consistent with the unified SABRE fit, the Hill slope *n* was released from unity and constrained to a single shared value across all groups.

## Appendix Estimation of Relative Efficacies Assuming Sigmoid Response and Occupancy Functions

Assuming straightforward hyperbolic functions (i.e., *n* = 1) for two compounds of different efficacy (*E*_max_) and potency (EC_50_), the effect *E* (response) depends on ligand concentration [L] as:

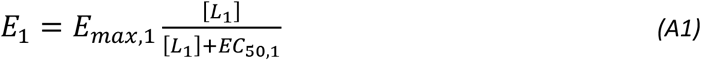

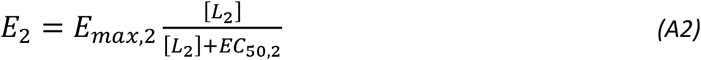

With this assumption, the ratio of ligand concentrations [L_1_] and [L_2_] causing the same effect *E*, which is small to ensure that is obtainable even with the weak partial agonists (see Figure below), can be obtained following the approach used to obtain the ratio of *equiactive molar ratios* by Ehlert and co-workers (EAMR; i.e., ratio of molar concentrations that produce the same response at low enough concentrations – later termed *intrinsic relative activity, RA*_i_) [23, 24]:

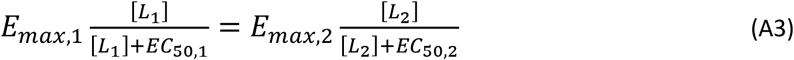

Rearranging to get the ratio of ligand concentrations:

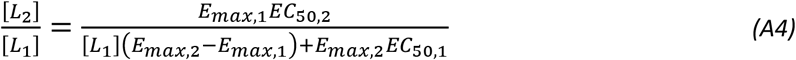

which at sufficiently low ligand concentrations approaches the EAMR [23]:

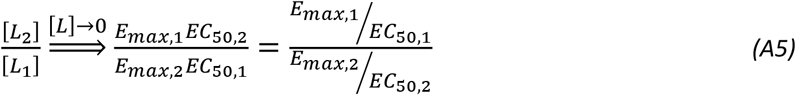

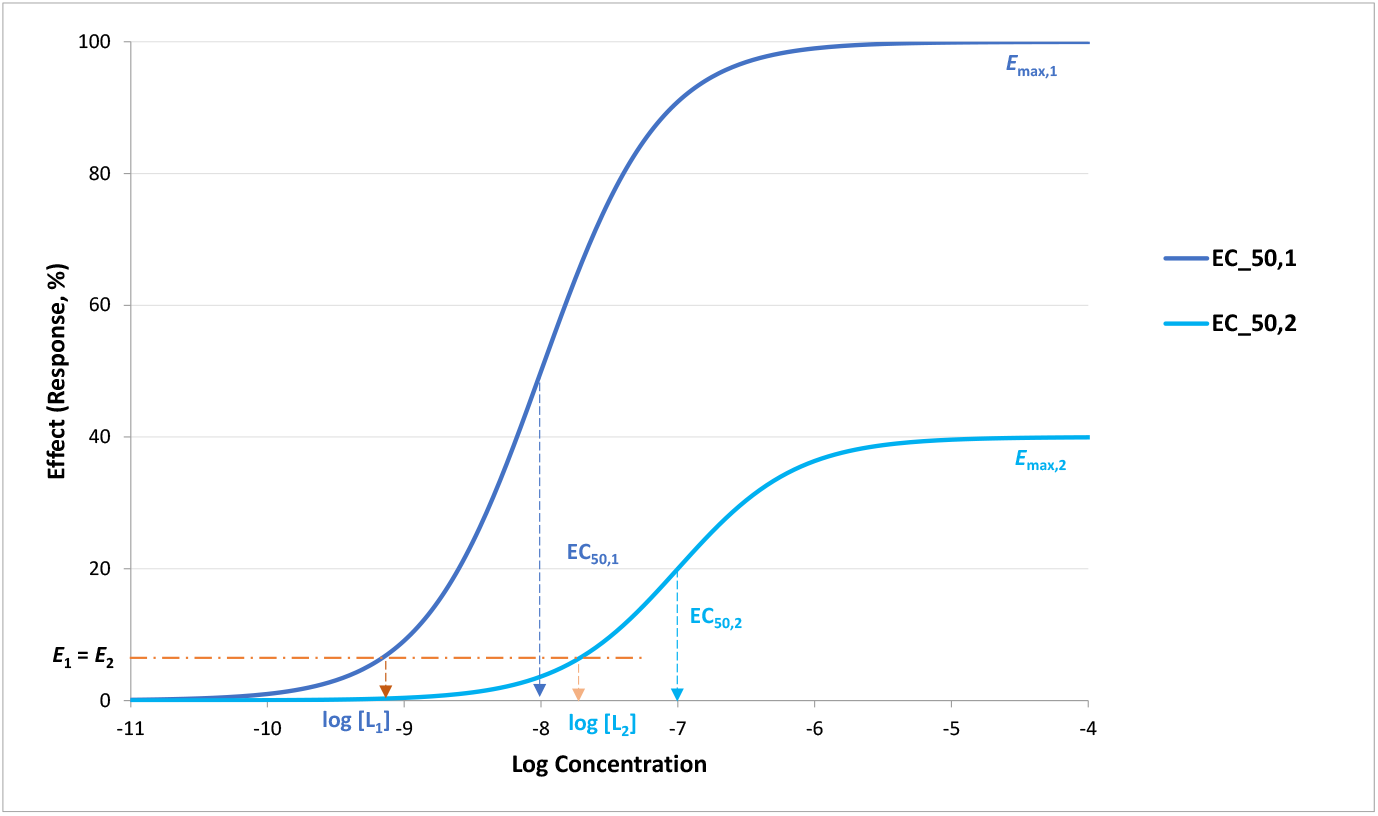

This *E*_max_/EC_50_ ratio is a frequently used basis of bias quantification, i.e., a logarithmic bias factor is obtained by calculating the shift ΔΔlog(*E*_max_/EC_50_), i.e., Δ_Pathway 2 vs 1_Δ_Compound Test vs Ref_ log(*E*_max_/EC_50_): (log(*E*_max,P₁,L_/EC_50,P₁,L_)–log(*E*_max,P₂,L_/EC_50,P₂,L_)) – ((log(*E*_max,P₁,Lref_/EC_50,P₁,Lref_)– log(*E*_max,P₂,Lref_/EC_50,P₂,Lref_)) [25-28]. Comparing the change on *E*_max_/EC_50_ is replaced with that of the “transduction coefficient” τ/*K*_D_ if the operational model is used or it can also be replaced with *εγ*/*K*_d_ if SABRE is used (assuming *n* = 1; see Appendix 2 of [1] and eq. 23 here).

To compare the relative efficacies of two compounds in the same assay fitted with sigmoid responses, one can follow the simple original approach by Furchgott, and assume that at conditions that produce equal responses, the ratio of efficacies (i.e., relative efficacies) is the inverse of the ratio of occupied receptors producing it [5]. Thus, with the notation used by Furchgott:

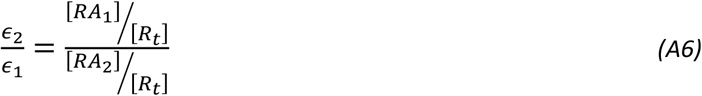

Or with the notation used for SABRE:

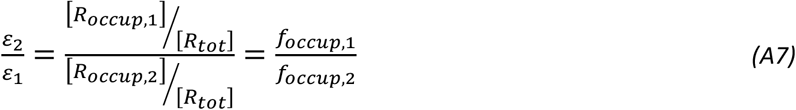

If one assumes that receptor occupancy follows a classic law of mass action and, hence, a sigmoid response function (on log scale) characterized by the *K*_d_ dissociation constant (which is determined here from responses measured at different receptor levels and fitted with individual functions, see eq. 14), *f*_occup_ for compound *i* is described by:

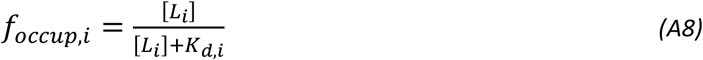

Thus, the ratio of occupancies as a function of ligand concentrations is

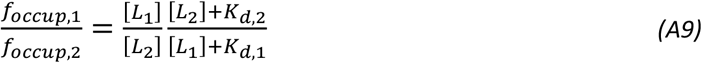

At low enough concentrations, which are assumed here in order to have low enough effects (see Figure and eq. A5), [L_1_] << *K*_d,1_ and [L_2_] << *K*_d,2_; thus, this ratio becomes:

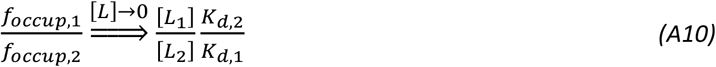

Using this in the equation for relative efficacies derived above (A7) and then introducing the ligand concentrations that produce the same (small) effect from eq. A5:

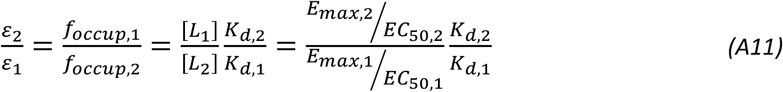

Which can be rearranged as

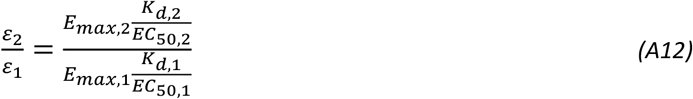

Therefore, comparing relative efficacies can be done by comparing *E*_max_,*K*_d_/EC_50_ values. The same relationship (eq. A12, eq. 15) can be obtained much more easily from the SABRE approach by using eq. 26, i.e.,

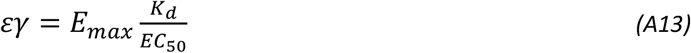

to express the ratio of efficacies (*ε*) and noticing that *γ* is eliminated as the pathway and, hence, the pathway amplifications are the same.

## Notes

### Competing Interest Statement

The authors have declared no competing interest.

### Summary of Updates

Added Appendix 1 with derivation of formula for relative efficacies assuming sigmoid response and occupancy functions. Updated manuscript text to account for this plus include a brief discussion of its use.

